# Motor cortex signals corresponding to the two arms are shared across hemispheres, mixed among neurons, yet partitioned within the population response

**DOI:** 10.1101/552257

**Authors:** K. Cora Ames, Mark M. Churchland

## Abstract

Primary motor cortex (M1) has lateralized outputs, yet M1 neurons can be active during movements of either arm. What is the nature and role of activity in the two hemispheres? When one arm moves, are the contralateral and ipsilateral cortices performing similar or different computations? When both hemispheres are active, how does the brain avoid moving the “wrong” arm? We recorded muscle and neural activity bilaterally while two male monkeys (*Macaca mulatta*) performed a cycling task with one or the other arm. Neurons in both hemispheres were active during movements of either arm. Yet response patterns were arm-dependent, raising two possibilities. First, the nature of neural signals may differ (e.g., be high versus low-level) depending on whether the ipsilateral or contralateral arm is used. Second, the same population-level signals may be present regardless of the arm being used, but be reflected differently at the individual-neuron level. The data supported this second hypothesis. Muscle activity could be predicted by neural activity in either hemisphere. More broadly, we failed to find signals unique to the hemisphere contralateral to the moving arm. Yet if the same signals are shared across hemispheres, how do they avoid impacting the wrong arm? We found that activity related to the two arms occupied distinct, orthogonal subspaces of population activity. As a consequence, a linear decode of contralateral muscle activity naturally ignored signals related to the ipsilateral arm. Thus, information regarding the two arms is shared across hemispheres and neurons, but partitioned at the population level.

## Introduction

The outputs of motor cortex (M1) are lateralized: most spinal projections influence the contralateral musculature. M1 lesions thus produce contralateral motor deficits (Liu and Rouiller, 1999; Murata et al., 2008; Passingham et al., 1983; Vilensky and Gilman, 2002). Similarly, electrical microstimulation activates contralateral musculature (Kwan et al., 1978; Sessle and Wiesendanger, 1982). The degree to which computations within M1 are lateralized versus shared across hemispheres remains less clear. The corpus callosum interconnects M1 across hemispheres, yielding the potential for extensive cooperation (Gould et al., 1986; Jenny, 1979; Jones and Wise, 1977). Callosally mediated interactions are readily revealed by paired-pulse TMS protocols and can involve net facilitation or suppression (Ferbert et al., 1992; Hanajima et al., 2001; Meyer et al., 1995). An obvious role for inter-hemispheric cooperation is coordination of bimanual movement (Donchin et al., 1998; Kermadi et al., 1998). Yet there is evidence that unimanual movements also involve sharing of information across hemispheres.

Most physiological studies of unimanual movements have focused on activity contralateral to the moving limb, on the grounds that contralateral activity is most functionally relevant and likely to be most prevalent. Yet studies investigating ipsilateral activity have found that it can be robust. Ipsilateral activity is minimal for tasks performed primarily with the digits (Matsunami and Hamada, 1981; Tanji et al., 1988; Aizawa et al., 1990) but prevalent during movements of the upper arm, such as reaching to remove food from a drawer (Kermadi et al., 1998; Kazennikov et al., 1999), or performing center-out reaching movements (Donchin et al., 2002; Steinberg et al., 2002; Cisek et al., 2003; Ganguly et al., 2009).

While the presence of ipsilateral activity is established, the nature of that activity is less clear. Few studies have directly compared neural response patterns when the same movement is performed by one arm versus the other. In premotor areas, delay-period responses can encode information about an upcoming reach (Cisek et al., 2003) or grasp (Michaels and Scherberger, 2018) independently of which arm would subsequently move, suggesting that preparatory activity is largely effector independent. However, activity during movement was more effector-dependent, for both premotor cortex and M1 (Cisek et al., 2003). Steinberg and colleagues (2002) reported similar single-neuron directional tuning regardless of which arm was moving, yet also found evidence for effector-dependent population-level encoding of direction.

If responses are effector-independent (i.e. similar regardless of the moving arm) then the relationship between hemispheres is necessarily simple: both contain the same information, encoded in the same manner. In contrast, if there exist strongly effector-dependent responses, that would raise additional questions. Are ‘lower-level’ signals (e.g., those describing muscle activity) more prevalent in the contralateral hemisphere? More generally, which signals are shared across hemispheres? How is neural activity structured such that only one arm moves even if both hemispheres are active?

We investigated these questions using a novel ‘cycling’ task, performed with either the left or right arm. We recorded neural activity from both hemispheres simultaneously. In separate sessions we recorded muscle activity bilaterally. Single neurons responded robustly regardless of which arm performed the task. Yet responses were strongly effector-dependent: for a given neuron, response patterns pertaining to the two arms were essentially unrelated. Despite profoundly effector-dependent single-neuron responses, we found no evidence that certain signals were present in one hemisphere but not the other. For example, muscle activity could be decoded equally well from contralateral or ipsilateral neural activity. More broadly, the population response across hemispheres appeared isomorphic; any signal present in one could also be found in the other. Thus, activity in a given hemisphere contains similar information during movement of one arm versus the other, yet that information is distributed very differently across single neurons. This might appear to yield a paradox: how can M1 be robustly active without driving the contralateral arm? A solution emerged when we examined the correlation structure between neurons, which changed dramatically depending on which arm was used. As a result, arm-specific signals were partitioned into orthogonal dimensions, allowing a simple decoder to naturally separate signals related to the two arms.

## Materials and Methods

### Terminology

We adopt the following terminology. For neurons in a given hemisphere, we refer to the contralateral arm as the “driven arm” (reflecting the strong connections to the contralateral spinal cord). We refer to the ipsilateral arm as the “non-driven arm.” Thus, for a neuron recorded from the right hemisphere, the left arm is the driven arm and the right arm is the non-driven arm. For the muscles, the driven arm is the arm upon which the muscle acts. Similarly, for a given arm, the contralateral cortex is referred to as the “driving cortex,” while the ipsilateral cortex is referred to as the “non-driving cortex.”

### Behavior

All animal procedures were approved by the Columbia University Institutional Animal Care and Use Committee. Data were collected from two male monkeys (*Macaca mulatta*) while they performed a cycling task for juice reward. Experiments were controlled and data collected under computer control (Real-time Target Machine: Speedgoat, Liebfeld, Switzerland). While performing the task, each monkey sat in a custom primate chair with the head restrained via surgical implant. A screen displayed a virtual environment through which the monkey moved. The monkey grasped a custom pedal with each hand, with the hands lightly restrained with tape to keep them in a consistent position on the pedals. The pedal itself was also designed to encourage a consistent hand position, and included a handle and a brace that reduced wrist motion. Pedals turned a crank-shaft attached to a motor (Applied Motion Products, Watsonville, California, USA). A rotary encoder within the motor reported position with 1/8000 cycle precision. The motor used information regarding angular position and its derivatives to provide forces yielding virtual mass and viscosity.

Monkeys cycled the pedal to control their position in the virtual environment (Figure 1A). For a block of twenty consecutive trials, one arm was the “performing arm,” and the other was the “non-performing arm.” The angular position of the performing arm’s pedal was mapped directly to linear position in the virtual world. The non-performing arm’s pedal was required to remain within a window (± 0.05 and ± 0.07 cycles for monkey E and F) centered at the bottom of the cycle. Movement outside that window caused trial failure followed by a short time-out. Monkeys adopted a stereotyped position within the window and moved little from that position while cycling with the other pedal. The position range explored within a trial averaged 0.006 cycles (monkey E) and 0.029 cycles (monkey F). Following completion of each twenty-trial block, the next block was signaled by a 5-second period during which the motor delivered a gentle “buzzing” to the upcoming block’s performing arm. Blocks were presented in randomized order (Figure 1C). Monkeys also executed blocks of trials where both arms cycled together (bimanual task variant), which are not analyzed in this study.

**Figure 1:**
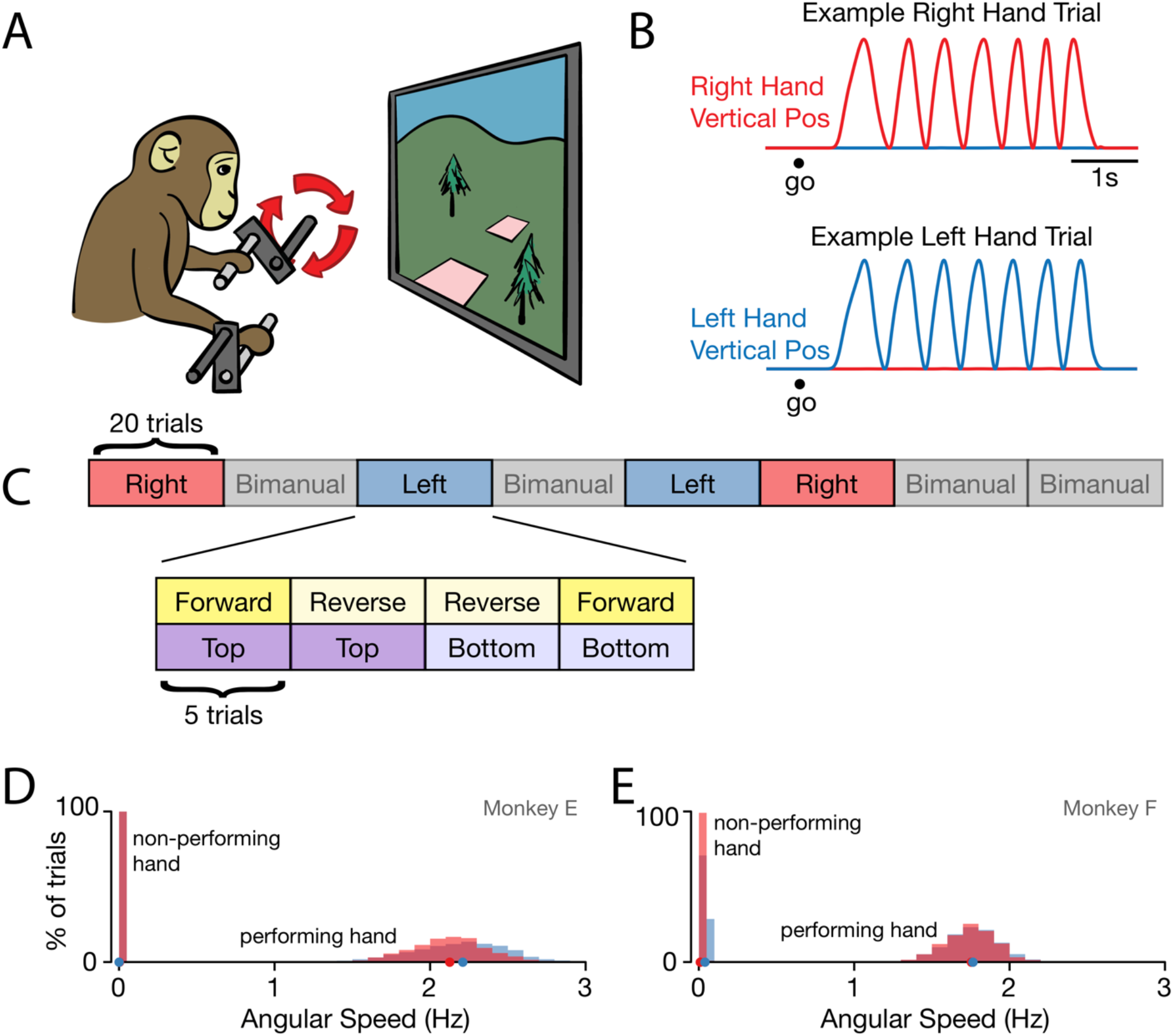
Behavior. (A) Task schematic. Cycling one of the two pedals produced progress through a virtual environment. The other pedal had to remain stationary. This schematic simplifies the physical setup. In particular, pedals employed a handle that ensured consistent hand posture and a brace that minimized wrist movement. (B) Behavior on two example trials. After a go cue, the monkey cycled for seven cycles with one hand while holding the other hand stationary. *Red trace*: Right hand vertical position. *Blue trace*: Left hand vertical position. (C) Task structure. Blocks of right hand, left hand, and bimanual conditions were presented in pseudorandom order. Within each block of 20 trials, trials were presented in sub-blocks of 5 trials for each combination of cycling direction and starting position. (D) Distributions of cycling speed for the performing and non-performing hands. The average angular speed was computed for each trial. Distributions are across trials. *Red*: Right hand. *Blue*: Left hand. Dots show distribution medians. Data are for Monkey E. (E) Same as D but for Monkey F.

During each trial, the monkey progressed from an initial target (a stationary white square on the ground) to a final target. The acceptance window was +/− 0.15 cycles for Monkey E and +/− 0.01 cycles for Monkey F. While stopped, the motor provided slight forces to Monkey F’s performing arm to enable the pedal to remain in this very stereotyped position with minimal muscle activation. While holding, the monkey was also required to maintain the performing arm below a speed threshold: 0.01 cycles/s for Monkey E and 0.0125 cycles/s for Monkey F. At the start of each trial, the initial target appeared one to two cycles in front of the monkey. The monkey cycled to and acquired this target. After 1000-2000 ms (monkey E) or 600-1000 ms (monkey F) the initial target disappeared and the final target appeared seven cycles ahead of the current position. The monkey cycled to this final target. Once the final target was acquired, the monkey remained still within the target to receive a juice reward.

Within each twenty-trial block there were four behavioral conditions. These conditions varied in the starting position of the pedal and the required cycling direction (Figure 1C). The initial target was located such that the pedal position necessary to acquire that target was either at the top (top-start) or at the bottom (bottom-start). The final target was a whole number of cycles away, and was thus acquired with the same pedal position. ‘Forward’ and ‘backward’ conditions differed in the cycling direction necessary to progress through the virtual environment. During forward cycling, the hand moved away from the body at the top (much as the foot does when pedaling a bicycle). During backward cycling, the hand moved towards the body at the top. Cycling direction was cued by the color of the landscape in the virtual world: green for forward, orange for backward. Each of the four combinations of starting position and cycling direction was performed in sub-blocks of 5 trials. Sub-block order was identical for each block (Figure 1C).

### Surgical Procedures and neural recording

Monkeys were anesthetized and a headcap was implanted under sterile conditions. A 19-mm diameter cylinder (Crist Instruments) was placed above the primary motor cortex of each hemisphere, guided by structural MRI performed prior to surgery. The skull remained intact under the cylinder, covered with a thin layer of dental acrylic. Prior to recording, monkeys were anesthetized and a 3.5-mm diameter burr hole was drilled by hand through the dental acrylic and skull, leaving the dura intact. Over the course of the experiment, multiple burr holes were opened at different locations. Following recording, burr holes were closed with dental acrylic, allowing the skull to heal.

After opening a burr hole, we first recorded neural activity using conventional single electrodes (Frederick Haer Company) to assess whether neurons in that location were task-modulated. We performed intracortical microstimulation and muscle palpations to confirm that recordings were within the arm region of M1. We then recorded neural activity with 24-channel V-Probes (Monkey E), or 32-channel S-Probes (Monkey F) (Plexon Inc, Dallas, Texas, USA). We lowered one probe into each hemisphere each day, removing the probe at the end of that session. Probes were moved to different locations within each burr hole on each recording day. Neural signals were processed and recorded using a Digital Hub and 128-channel Neural Signal Processor (Blackrock Microsystems, Salt Lake City, Utah, USA). Threshold crossings from each channel were recorded and spike-sorted offline (Plexon Offline Sorter). Unit isolation was assessed based on separation of waveforms in PCA-space, inter-spike interval histograms, and waveform stability over the course of the session. Analyses consider stable, well-isolated single and multi-unit isolations. Multi-unit isolations consisted of identifiable spikes (i.e., not merely threshold crossings) from two (or occasionally more) neurons that could not be distinguished with confidence. All example firing rates shown in figures are from single units. We recorded 263 units in the left hemisphere and 270 units in the right hemisphere for monkey E, and 338 units in the left hemisphere and 279 units in the right hemisphere for monkey F.

### EMG recording

On a separate set of days from the neural recordings, we recorded intramuscular EMG signals from the following muscles: Biceps brachii (long and short head), triceps brachii (medial, long, and lateral heads), deltoid (anterior, lateral, and posterior head), latissimus dorsi, pectoralis, trapezius, and brachioradialis. Pairs of hook-wire electrodes were inserted ∼1cm into the belly of the muscle being recorded at the beginning of each session and removed at the end of the session. On each session, 1-3 EMG recordings were made per arm. Electrode voltages were amplified, bandpass filtered (10-500 Hz) and digitized at 1000 Hz (Monkey E) or 30000 Hz (Monkey F). Recordings were not considered further if they contained significant movement artifact or weak signals. Offline, EMG records were high-pass filtered at 40 Hz, rectified, and smoothed with a 25-ms Gaussian. This produced a measure of intensity versus time, which was then averaged across trials.

### Trial Averaged Firing Rates

The spike times of each neuron on each trial were converted to a firing-rate by convolving spikes with a 25-ms Gaussian. To produce trial-averaged firing rates, we first aligned all trials on a common time: the moment when the first half-cycle was completed. This nicely aligned behavior across trials during the first cycle of each trial. However, because trials lasted multiple seconds, small differences in cycling speed could accumulate and cause considerable misalignment of behavior across trials (Figure 2A). The resulting misalignment of spikes (Figure 2B) would, if averaged without further alignment, yield an unrepresentative average firing rate (the same problem would impact averages of muscle activity and kinematic variables). Thus, the time-base on each trial was adjusted such that each cycle lasted 500 ms (excluding the first and last half-cycles), matching the typical 2 Hz cycling speed (Figure 2C). This procedure altered the time-base of individual trials very modestly, yet maintained appropriate alignment across trials (Figure 2D), and produced trial-averaged estimates of the firing rate (Figure 2E) that are representative of what occurred on single trials.

**Figure 2:**
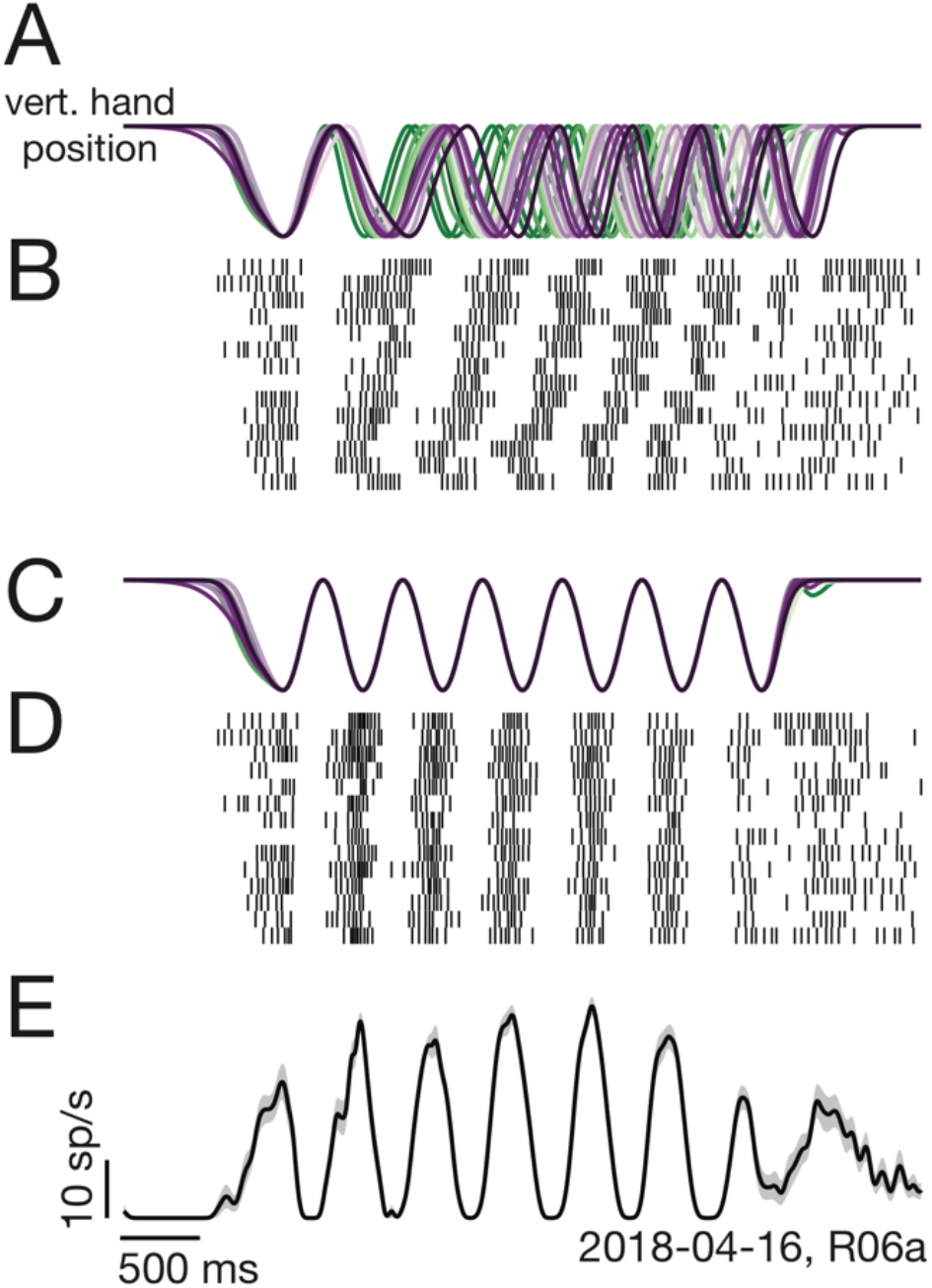
Two-step trial alignment procedure. (A) Vertical hand position on all trials (within one session) for an example condition. Data are aligned to the first half-cycle of movement. Trials are colored *green* to *purple* based on the average cycling speed for that trial. (B) Raster plot of spike times for an example neuron, for the same trials as in A. Trials are ordered by average cycling speed and aligned as in A. (C) Hand position traces after the second alignment step: adjusting the time-base of each trial so that cycling during the middle 6 cycles matched the typical 2-hz pedaling speed. (E) Spike times, as in B, after the second alignment step. (F) Average firing rate calculated after the second alignment step. *Black*: Mean firing rate. Gray shading: standard error across trials.

Figure 2D shows spike rasters to illustrate improved alignment of neural activity. However, we stress that spikes were always converted to rates before modification of the time-base. Thus, the alignment procedure did not alter the values of the estimate of firing rate; it simply slightly modified when those values occurred.

Most analyses of firing rates employed the middle cycles (2-5), excluding the first cycle and the last two cycles. This focused analysis on the steady-state response, rather than on responses associated with starting, stopping, or holding. This aided interpretation in two ways. First, muscle activity in the non-performing arm was particularly weak during middle cycles (in contrast, modest activity was occasionally observed when stopping). Focusing on middle cycles largely sidesteps concerns that activity ipsilateral to the performing arm is related to muscle activity in the non-performing arm. Second, we wished to focus key analyses on the rhythmic pattern of firing rate modulation, rather than on overall changes in net firing rate when moving versus not moving. As one example, when predicting muscle activity from neural activity, it is relatively ‘easy’ to capture the generally elevated activity level during movement, resulting in high *R*^*2*^ values even if predictions fail to capture cycle-by-cycle activity patterns. We wished to avoid this, and to consider predictions successful only if they accounted for rhythmic response aspects.

### Single-neuron analyses

We wished to compare, for each neuron, the strength of modulation when the driven versus non-driven arm performed the task. By modulation, we mean the degree to which a neuron’s firing rate varied within cycles, between cycles, and/or between conditions (forward versus backward, and top-start versus bottom-start). We compiled a single firing rate vector, *r*_*driven*_, concatenating the firing rate vectors across the four conditions where the driven arm performed the task. *r*_*driven*_ was thus of size *ct* where *c* is the number of conditions and *t* is the number of times during the middle cycles of one condition. We defined *Modulation*_*driven*_ as the standard deviation of *r*_*driven*_, which captures the degree to which the average firing rate varies across time and condition. *Modulation*_*non-driven*_ was computed analogously.

To assess the degree to which a neuron was more strongly modulated when the driven versus non-driven arm performed the task, we computed an arm preference index:

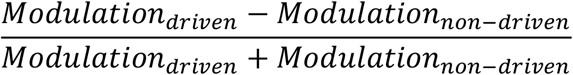

This arm preference index is zero if a neuron is equally modulated regardless of the arm used, approaches one if modulation is much larger when using the driven arm, and approaches negative one if modulation is much larger when using the non-driven arm.

### Firing-rate impact of small movements of the non-performing arm

We wished to control for the possibility that neural responses, when the non-driven arm performs the task, might be related to small movements of the non-performing arm. For each trial, we computed the mean (absolute) speed of the non-performing arm. For each condition, we divided trials into those with speeds greater versus slower than the median. We did not apply this analysis if there were fewer than 8 trials for that condition. This could occur if a neuron was well isolated for only part of a recording session.

After dividing, we recomputed the mean firing rate for each of the two pools of trials, yielding one firing rate when the non-performing arm moved modestly, and another when it was virtually stationary. For each timepoint, we asked whether these two firing rates were more different than expected given sampling error. This was accomplished via a bootstrap in which trials were divided randomly, rather than based on speed. We performed 1000 such random divisions. Differences were considered significant if they were larger than for 95% of the random divisions.

### Normalization

Because the absolute voltages of EMG traces are largely arbitrary, the scale of muscle activity could be quite different for different muscles. The response of each muscle was therefore normalized by its range. Neural responses were left un-normalized for single-neuron level analyses. However, for population-level analyses, responses were normalized to prevent results from being overly biased toward the properties of a few high-rate neurons. For example, principal component analysis seeks to capture maximum variance, and a neuron with a firing rate modulation of 100 spikes/s would contribute 25 times as much variance as a neuron with a modulation of 20 spikes/s. Reducing that discrepancy encourages principal component analysis to summarize the response of all neurons. For these reasons, we normalized the firing rate of each neuron, using the equation 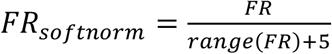. The addition of 5 to the denominator produces ‘soft’ normalization, and ensures that we don’t magnify the activity of very low-rate neurons. We have used this value previously (e.g., Lara et al. 2018 eLife; Russo et al. 2018 Neuron) as it strikes a reasonable balance between focusing analysis on all neurons, while still allowing high firing-rate neurons to contribute somewhat more than very low-rate neurons.

### Population Predictions

To predict muscle activity from neural activity, we used Partial Least Squares (PLS) regression (plsregress in MATLAB). For each set of neurons *X* and muscles *Y*, PLS regression finds matrices *W, V*, that maximize the covariance between *XW* and *YV*, under the constraint that *W, V* are of rank *r* (which must be specified). PLS is similar to Canonical Correlation Analysis, in that it seeks linear transformations of the data that maximize similarity. However, Canonical Correlation Analysis maximizes correlation, and can therefore often be biased toward small dimensions which are coincidentally well-correlated. In contrast, PLS regression maximizes covariance and thus seeks correlated signals that are also high variance. Once *W* is found, *Y* is predicted from *XW* via standard linear regression. Employing *XW* (which has only *r* columns of regressors) rather than *X* (which has hundreds of columns) greatly reduces overfitting. An advantage of PLS regression is that the regularized solution respects not only the correlations in *X* (as for PCA regression) but also the correlations in *Y*.

All predictions involved the middle cycles (2-5) of movement. We first picked one behavioral condition (e.g. top-start, forward, right hand performing) as our test condition. We then set the training condition to be the behavior with the same cycling direction but the opposite starting pedal position (e.g. bottom-start, forward, right hand performing). We first mean-centered the data, such that average neural and muscle activity was zero for each condition. We then ran PLS regression on the training condition to find a rank-*r* matrix *B* that predicts muscle activity from neural activity. To select the optimal rank, we selected one cycle from our test condition to serve as validation data. We assessed performance on this validation cycle and selected the rank, *r*\*, and the corresponding weight matrix, *B*\*, that generated the maximal validation *R*^*2*^. We assessed prediction performance (generalization) on the remaining cycles of our test data. This procedure was repeated for each behavior and hemisphere. Generalization performance was assessed based on population percent variance explained. We considered *Y*_*pred*_ = *XB*\*, and computed 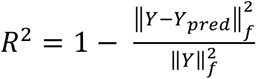, where ‖.‖_*f*_ indicates the Frobenius norm.

We used a similar procedure to assess how well neural activity could be predicted from neurons in the same or opposite hemisphere. We randomly divided driving-cortex neurons into two halves: *X*_*driving*_ and *Y*_*driving*_. We also considered a random subpopulation of non-driving cortex neurons, *X*_*non-driving*_, selected to have the same number of columns (neurons) as *X*_*driving*_. Using PLS regression, we calculated generalization performance, 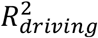, when predicting *Y*_*driving*_ from *X*_*driving*_. We similarly computed 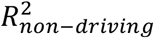 when predicting *Y*_*driving*_ from *X*_*non-driving*_. We used the same train/validate/test procedure described above, assessing generalization performance on a held-out condition. We repeated this process with 125 different random divisions per condition, yielding a total of 1000 values of 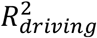 and 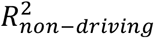 per monkey.

### Dimensionality Reduction

Dimensionality reduction was performed via principle components analysis (PCA). We typically ran PCA on neural data from a sub-set of behavioral conditions. We concatenated neurons’ soft-normalized FRs from the desired conditions to generate a data matrix A of size *(c* × *t, n)*, where c was the number of conditions, t was the number of timepoints included per condition, and n was the number of neurons. We applied PCA to A to obtain matrices X and V such that *X* = *AV*, where X is the projection of the data onto the principal components (PCs) and V contains the weights from neurons to PCs. To project other behavioral conditions into the same space, we could construct a new data matrix *A*′ using these conditions’ FRs. We then multiply by V, such that the new projections *X*′ are defined by *X*′= *A*′*V*.

### Trajectory Tangling

We assessed trajectory tangling as described in(Russo et al., 2018). To parallel the other analyses of population activity, trajectory tangling was computed for the middle cycles. Neural activity (or muscle activity) was reduced to the top eight dimensions using PCA. We then calculated tangling, *Q(t)*, at each time point:

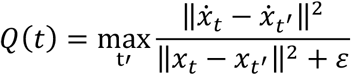

where *x*_*t*_ is the neural state at time *t*, 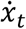 is the temporal derivative of the neural state, ‖.‖ is the Euclidean norm, and ε is a small constant that prevents division by zero (here, set to 10% of the total neural variance across the top eight dimensions). *Q(t)* is large if the neural state at time *t* is close to the neural state at a different time, but the two states have very different derivatives.

Trajectory tangling was computed across all times for a given set of conditions – e.g. all conditions where the right arm performed the task. Thus, *t* indexes across all times and conditions for one arm. Trajectory tangling was computed separately for each hemisphere, and for the muscle population in each arm. For a given quantity (e.g., muscle activity) the two distributions (one per arm) for that monkey were combined and the cumulative density was computed.

### Predicting non-performing arm EMG

Above we described our methodology for assessing how well neural activity in a given hemisphere predicts muscle activity in the arm performing the task. We used a similar methodology to address a related but different question: whether a unified linear decoder, based on activity across both hemispheres, can predict muscle activity both when that arm performs the task (and robust EMG needed to be decoded) and when the other arm performs the task (and near-zero EMG should be decoded). As above, we used PLS regression and focused on the middle cycles. For this analysis, we predict EMG activity using all neurons, regardless of hemisphere.

We first assessed generalization performance using a ‘train-moving’ decoder, which was trained using only conditions where the relevant arm performed the task, and was then asked to generalize to conditions where the arm did not move. This is a potentially challenging form of generalization, as the decoder must predict EMG activity in a situation (arm not moving) very different from the situation in which it was trained (arm moving). We also computed generalization performance of a ‘train-both’ decoder, trained using a set of conditions that included the relevant arm performing and not performing the task. Generalization was to left-out conditions of each type.

For the train-moving decoder, we used the following division of training, validation, and testing, conditions: *Training*: Direction 1, Start Position 1, Arm moving. *Validation*: Direction 1, Start Position 2, Arm moving (one cycle). *Testing*: Direction 1, Start Position 2, Other arm moving. For the train-both decoder, we used the following division of training, validation, and testing: *Training*: Direction 1, Start Position 1, Arm moving and Direction 1, Start Position 1, Other arm Moving. *Validation*: Direction 1, Start Position 2, Arm moving (one cycle). *Testing*: Direction 1, Start Position 2, Other arm moving.

## Results

### Behavior

Two monkeys (E and F) were trained on a cycling task that could be performed with either arm (Figure 1A). Left and right hands each grasped a pedal. Monkeys performed blocks of left-hand and right-hand trials. Cycling the correct pedal produced motion through the virtual environment. Success required that the non-performing arm be kept still. On each trial, monkeys cycled from one target to another, located seven cycles away (Figure 1B). Targets were positioned so that cycling started and ended either at the top of the cycle (‘top start’) or at the bottom of the cycle (‘bottom start’).

Each combination of starting position and cycling direction was performed for five consecutive trials. The order of the four combinations was consistent within each 20-trial block (Figure 1C). Monkeys performed an average of 29 and 21 trials per condition per day (monkey E and F, respectively). Monkeys cycled quickly, with a median angular speed of 2.2 Hz and 1.8 Hz (monkey E and F; Figure 1D,E). In contrast, the non-performing hand moved very little (leftmost distributions in Figure 1D,E). Mean angular speed for the non-performing arm was 0.0016 cycles/s and 0.024 cycles/s (monkey E and F).

### Neural and muscle responses

We examined the average firing rate of neurons recorded in each hemisphere of M1 (1150 total isolations across two hemispheres and both monkeys). Firing rates were computed after temporally aligning behavior across trials (Figure 2). Neural responses were typically rhythmic (Figure 3E-H), and could be nearly sinusoidal (Figure 3E) or could contain additional higher-frequency structure (Figure 3G). For comparison, we recorded the activity of the major muscles in both arms (48 total recordings). The temporal features of muscle responses (Figure 3A-D) in many ways resembled those of single-neuron responses. However, muscles and neurons were quite different in the degree to which responses were restricted to movements of a single arm.

**Figure 3:**
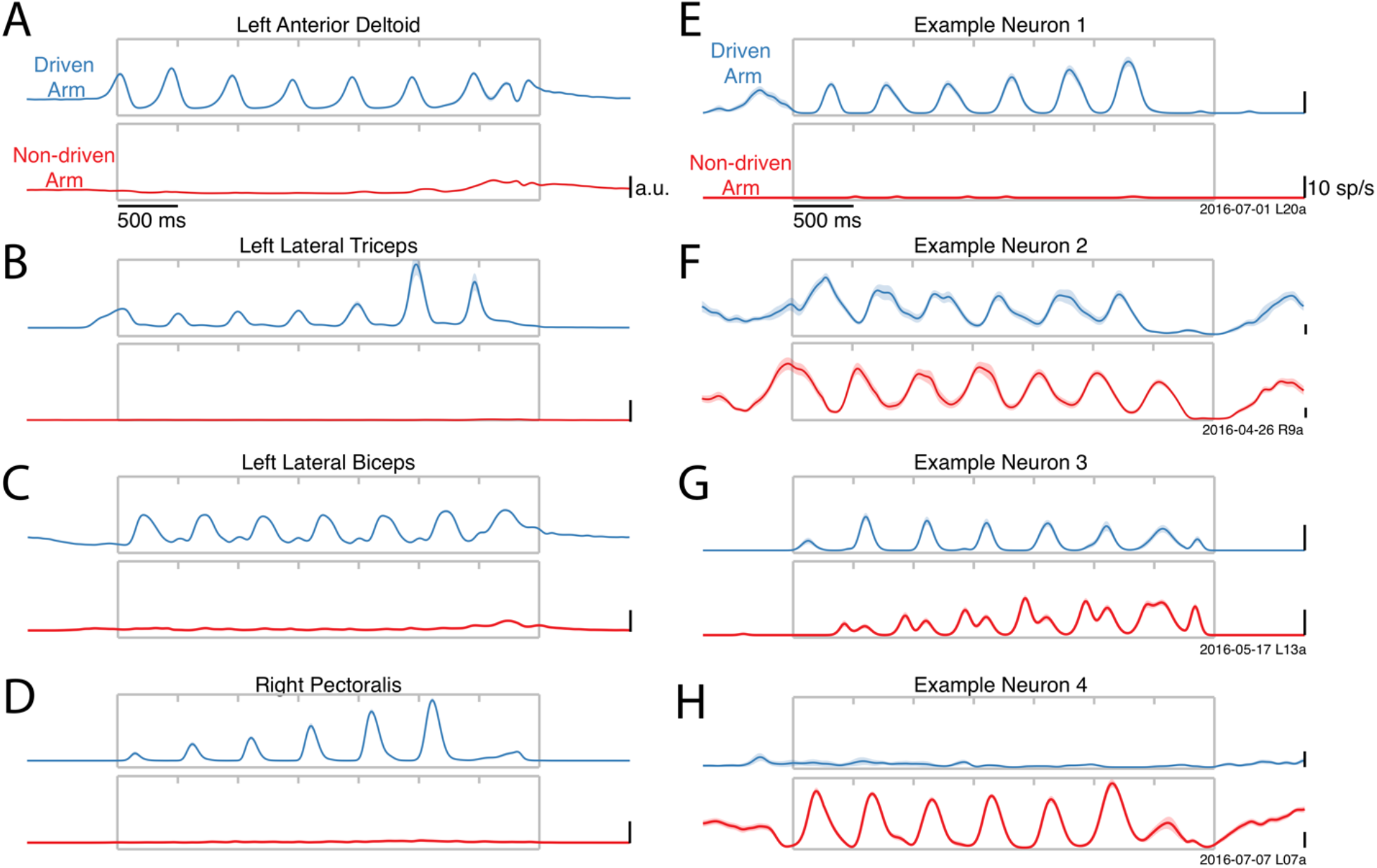
Activity versus time for four example muscles (A-D) and four example neurons (E-H). Each panel shows activity for one condition performed with either the driven arm (*blue*) or the non-driven arm (*red*). Each trace plots trial-averaged activity, with flanking envelopes (sometimes barely visible) showing standard errors. Gray boxes indicate when the pedal was moving, with tick marks dividing each cycle. All data are from Monkey E.

Muscles exhibited robust activity only when their driven arm performed the task (Figure 3A-D). E.g., the left anterior deltoid (Figure 3A) was active when the left arm performed the task (*blue*) but not when the right arm performed the task (*red*). While expected, this direct confirmation is important because of the possibility that muscles might have been active in ways that didn’t move the pedal (e.g., co-contraction). Such activity could potentially have been substantial, complicating interpretation of neural activity. A few muscles exhibited weak activity when the task was performed by their non-driven arm (Figure 3A,C). However, this typically occurred only at the end of movement, consistent with tensing to aid stability during stopping.

In contrast to the muscles, neurons were typically active throughout the movement, regardless of whether the task was performed with their driven or non-driven arm. A few neurons were active only when cycling with the driven arm (Figure 3E), and on rare occasions a neuron was active only when cycling with the non-driven arm (Figure 3H). However, most neurons were active in both situations (Figure 3F,G). Furthermore, neural responses could be quite different when cycling with the driven versus non-driven arm. Neural response patterns could change in both phase (Figure 3F) and structure (Figure 3G) depending on which arm performed the task.

### Single-neurons are active during movements of either arm

To quantify the arm preference of individual neurons, we compared firing-rate modulation when the task was performed with the driven versus non-driven arm. Modulation was assessed as the standard deviation of the firing rate across timepoints, which captures the degree to which activity evolves with time. Average modulation was computed once across all conditions where the driven arm performed the task, and again across all conditions where the non-driven arm performed the task. We analyzed only the middle cycles of movement (excluding the first cycle and the last two cycles). This allowed us to quantify the “steady state” performance of the neurons, without starting and stopping transients, and aided comparison with the muscles. We computed an ‘arm preference index’: the difference in modulation for the driven versus non-driven arm, divided by the sum. This index ranges from −1 to 1, with the extremes indicating complete preference for the non-driven and driven arms respectively. An arm preference index of zero indicates that a neuron was equally responsive regardless of the arm being used.

To establish a baseline for comparison, we computed the arm preference index for each muscle. Arm preference indices were typically high for the muscles, confirming that muscles were active primarily when the task was performed with their driven arm. A few muscles showed weak activation regardless of the arm being used, resulting in lower indices. However, most muscles had robust responses, and were much more active when the task was performed with the driven arm. For both monkeys, the median arm preference index was near unity (Figure 4A E: 0.86; F: 0.98; *blue dots*) and the modal response occurred at unity.

**Figure 4:**
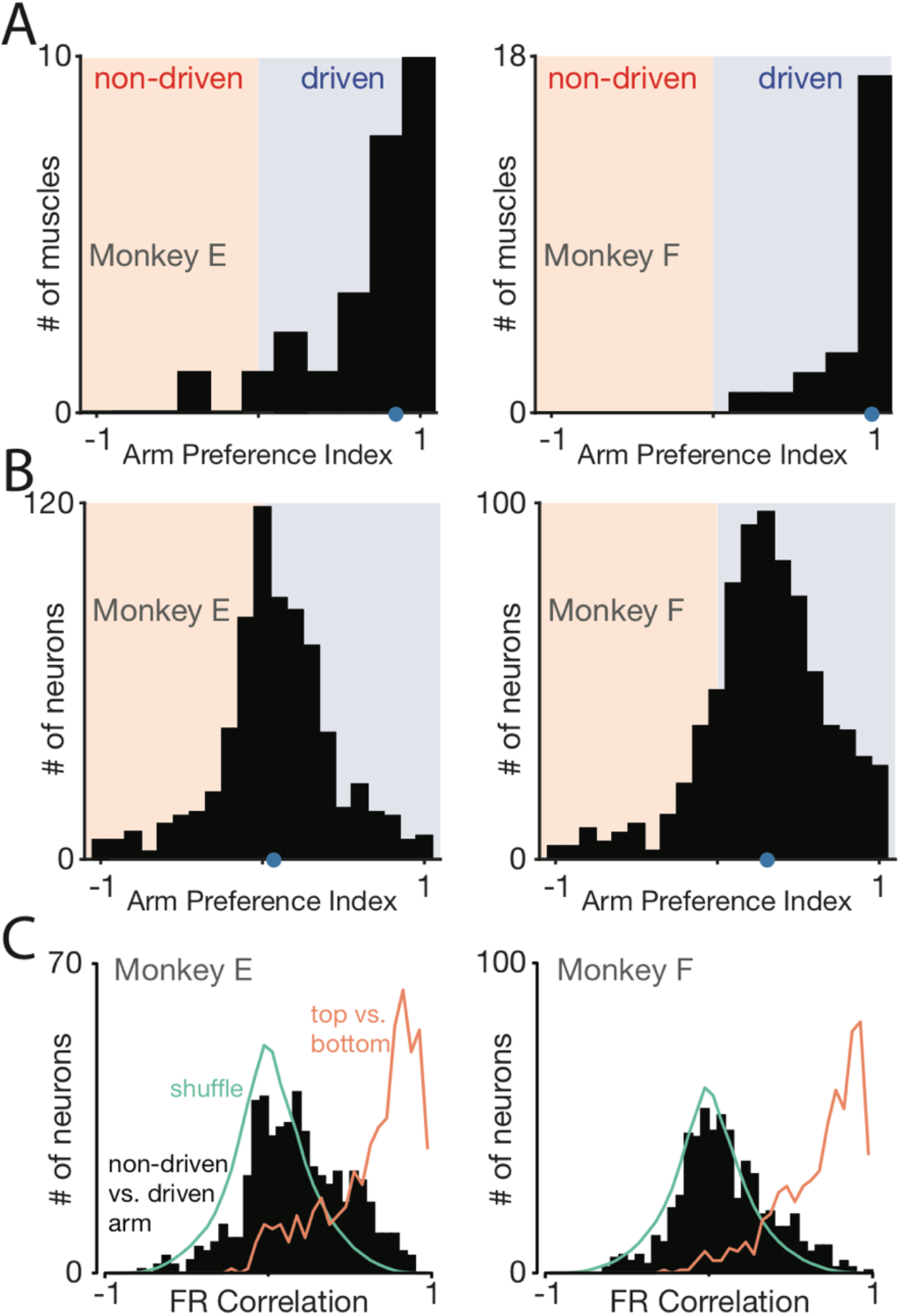
Muscle responses are lateralized while neural responses are not. (A) Histograms of arm preference index for all recorded muscles. Shaded regions indicate preference for the non-driven arm (*red*) and driven arm (*blue*). *Blue dot* indicates median arm preference. (B) Histograms of arm preference index for all recorded neurons. (C) Histograms summarizing, for single neurons, similarity of responses when the task is performed with the driven versus non-driven arm. For each neuron, we computed the correlation between those two responses. *Black* histograms plot the distribution of such correlations across all neurons. *Green trace*: expected distribution if there is no relationship between responses corresponding to the two arms. This was computed via a shuffle procedure. *Orange trace*: control demonstrating that high correlations are observed, as expected, when comparing responses during top-start versus bottom-start conditions.

In contrast, neurons rarely had arm preference indices near unity (Figure 4B). Instead, the distribution of arm preference indices was centered slightly above zero (median = 0.07 and 0.31 for the two monkeys). Thus, neural responses were much more likely than muscle responses to be similar in magnitude regardless of which arm performed the task. Furthermore, many neurons had arm preference indices < 0, indicating stronger modulation when the non-driven arm performed the task (Monkey E: 201/533 neurons; Monkey F: 107/617 neurons).

Thus, neurons can be quite active even when the task is performed with their non-driven arm. Might such responses be related to small movements of the driven arm? This explanation is unlikely *a priori*. As described above, movements of the non-performing arm were small (Figure 1D,E) and corresponding muscle activity was weak (Figure 4A). In principle, neural responses related to weak muscle activity might be magnified via normalization or some other non-linearity. However, such magnification would need to be very strong. To match the median neural arm preference indices, muscle activity in the non-performing arm would need to be magnified by a factor of 12 (monkey E) and 52 (monkey F). Furthermore, magnification cannot account for the finding that neurons commonly had negative arm preference indices, while muscles rarely (monkey E) or never (monkey F) did.

We performed an additional control to ask whether neural responses, when the non-driven arm performed the task, were influenced by small movements of the driven arm. Such movements varied across trials (Figure 5A), allowing us to divide trials into those with movements larger than the median (very modest movement, *red*) versus lower than the median (nearly stationary, *black*). Average firing rates were very similar in these two cases, as illustrated for one example neuron in Figure 5B. Differences were significant at only a few moments (*black dots*, p < 0.05 via bootstrap across 1000 resamples). Those differences were small, and occurred at roughly the rate (5%) expected by chance. Across all neurons, 5% of data-points showed significant differences (Figure 5C,D). This equals the percentage expected by chance, and is thus consistent with no reliable impact of small movements. Applying this same analysis to the (weak) muscle activity in the non-moving arm revealed significant differences at double the chance rate (10% of data-points) and peaking at four times the chance rate near the end of the movement (21% of data-points).

**Figure 5:**
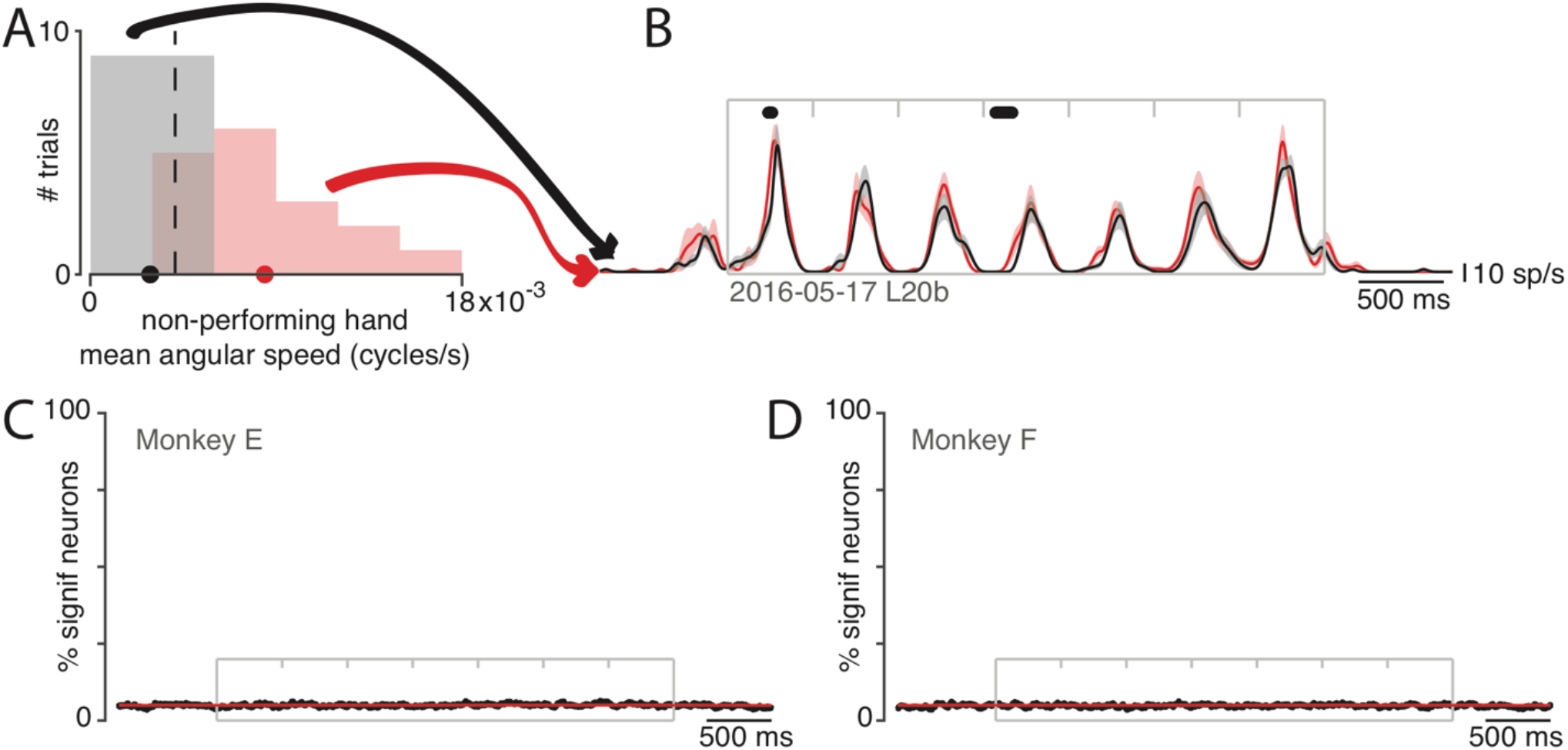
Small movements of the non-performing arm cannot explain modulation of neural activity within the non-driving cortex. (A) Analysis employed the distribution (across trials) of the mean angular speed of the non-performing arm. This distribution is shown for one condition, recorded on one day. Trials were divided into those with mean speed less than (*gray*) or greater than (*red*) the median (*vertical dashed line*). (B) Firing rate of one example neuron for these two groups: trials with speeds less than (*black*) and greater than (*red*) the median. Envelopes show standard errors of the mean. *Black dots* at top indicate times when the two rates were significantly different (p<0.05). Plotting conventions as in Fig. 3. (C) Percentage of neurons (*black trace*) showing a statistically significant difference (p<0.05) in firing rate for trials with speeds less than versus greater than the median. Differences occurred roughly as often as expected by chance (*red line* at 5%). *Gray box* denotes the time of movement. Each tick mark delineates a cycle. Data are for monkey E. Analysis is based on 426 neurons. (D) As in C but for Monkey F. Analysis is based on 479 neurons.

In summary, muscles were silent or at most weakly active when the task was performed with their non-driven arm. The weak activity that was present was statistically coupled to small movements of the driven arm. In contrast, neural responses were typically robust during movements of the non-driven arm, were present throughout the movement (not just at the end) and were not statistically linked to small movements of the driven arm. Prior studies have found that neurons can be active when a task is performed with their non-driven arm, although to varying degrees (Cisek et al., 2003; Donchin et al., 1998; Kermadi et al., 1998; Tanji et al., 1988). The present findings replicate the finding of weakly lateralized responses in motor cortex, and largely rule out potential explanations based on small residual movements of the driven arm.

### Neural response patterns are limb-dependent

One plausible explanation for weakly lateralized responses is that neural activity encodes higher-level, limb-independent features of movement. For example, activity might encode hand velocity, movement goal, or some other quantity, regardless of which limb is moving. Preparatory activity in the more anterior rostral premotor cortex (Cisek et al., 2003) can exhibit largely limb-independent responses. Might this also be true in motor cortex during movement? The two arms performed very similar movements in our task. Thus, limb-independence should be reflected by similar neural responses regardless of the performing arm. Instead, neural responses were strongly limb-dependent. Responses often differed in phase (Figure 3F) and/or structure (Figure 3G) depending on which arm performed the task.

To provide quantification, for each neuron we computed the correlation between the firing rate patterns when the driven versus non-driven arm performed the task (Figure 4C). Analysis considered only the middle cycles (2-5). This ensured that high correlations indicate similar response patterns, not simply firing rates that rise non-specifically during movement. On average the correlation was near zero (median correlation: 0.16 and 0.08 for Monkey E and F). Strongly correlated responses were very much the minority: only 18/533 (E) and 9/617 (F) neurons had correlations >= 0.75. Thus, for a given neuron, there was remarkably little relationship between responses when the task was performed with the driven versus non-driven arm. We used a shuffle manipulation to estimate the distribution of correlations if there were in fact no relationship. Each neuron’s response was matched with that of another random neuron, yielding a distribution of correlations expected by chance given the range of response patterns present in the data (*green*). The empirical distribution (*black*) was only modestly more positive than the chance distribution.

Might correlations appear artificially low if responses are weak or noisy? While sampling error will inevitably reduce correlations, this is unlikely to be the source of the low correlations we observed. Cycling evoked particularly strong neural responses with correspondingly small standard errors of the mean firing rate (Figure 3E-H, envelopes show SEM). We further addressed this concern by computing, for each neuron, the correlation between the firing rate for the top-start versus bottom-start conditions. Behavior was very similar for these two conditions during the middle cycles (after aligning phase), and correlations should thus be high. This was indeed the case (Figure 4C, orange distributions), confirming that sampling error did not impede the ability to measure high correlations.

These results rule out the hypothesis of a representation that is predominantly effector-independent. Individual-neurons showed very different responses depending on which arm performed the task – almost as different as if there were no relationship between the activity patterns associated with the two arms.

### Correlations between neurons are limb-dependent

Consider two neurons that have similar response patterns when the task is performed with the driven arm (Figure 6A-D plots four such example pairs). What occurs when the task is performed with the non-driven arm? From the analysis above, we know that the response pattern of each neuron will change. Does this occur in a coordinated fashion, such that the two neurons remain correlated with one another? This would be consistent with the idea that neurons with related responses ‘encode’ related features, and continue to do so in new contexts. In fact, this property was rarely observed. We did occasionally observe neurons that were strongly correlated when the driven arm performed the task, and remained strongly correlated when the non-driven arm performed the task (Figure 6A). Yet it was also common for correlations to invert (Figure 6B), for strong correlations to disappear (Figure 6C), or for neurons to undergo very different changes in response magnitude (Figure 6D).

**Figure 6:**
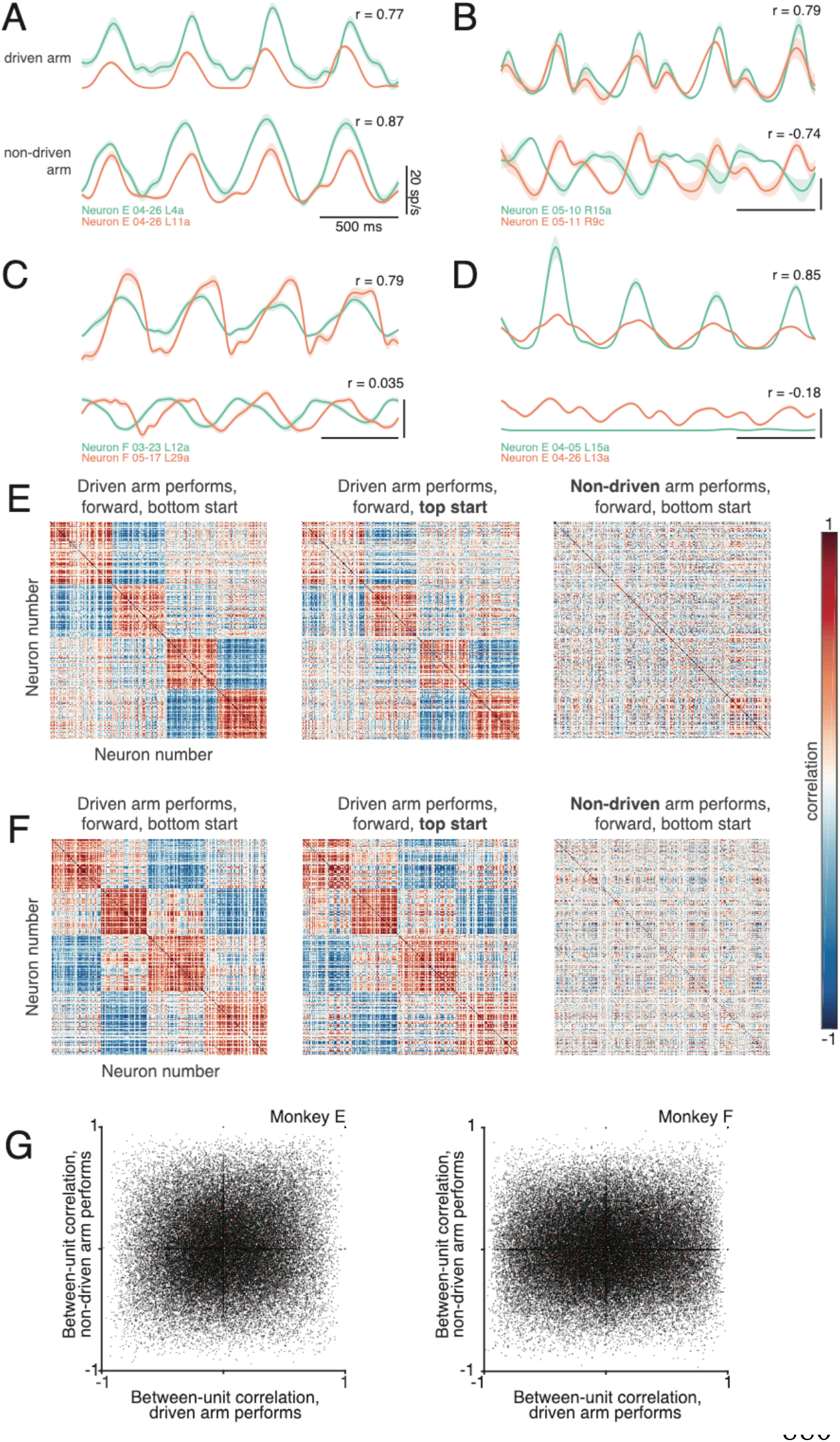
Correlations between neurons depend on which arm is used. (A) Average firing rates of two neurons (*green* and *orange traces*) during one condition, performed with the driven arm (top pair of traces) and non-driven arm (bottom pair of traces). Responses are shown for the middle cycles, which form the basis of the analysis below. For this example, the firing rates of the two neurons were strongly correlated when the driven arm performed the task, and remained so when the non-driven arm performed the task. Flanking envelopes show standard errors of the mean. (B-D) Responses of three other pairs of neurons. All pairs exhibit correlated responses when the task was performed with the driven arm. Correlations disappear or even invert when the task is performed with the non-driven arm. (E) Pairwise correlations between all neurons recorded from the left hemisphere of monkey E. Each panel plots the correlation matrix for one condition, indicated at top. Neuron ordering was based on data in the left column, and is preserved across columns. (F) same but for Monkey F. (G) Scatterplots of pairwise correlations. Each dot corresponds to a pair of units for a given condition, and plots the firing rate correlation when using the non-driven arm versus that when using the driven arm. To aid visualization, a randomly-selected 10% of data points are shown for this subplot.

We computed correlation matrices to quantify such effects across the population. To aid visualization, we ordered neurons to group responses that were similar when the task was performed with the driven arm, resulting in a block structure (Figure 6E-F, *left*). We asked whether this correlation structure remained similar when the task was performed with the non-driven arm. Note that it is possible for the correlation matrix to remain identical, even if every neuron changes its response, so long as correlated neurons remain correlated. Instead, the correlation structure was dramatically altered. As a result, the original ordering no longer groups neurons with similar response properties (*right column* of Figure 6E-F).

This change in correlation structure was not due to correlations being largely spurious, as could occur if estimated firing rates were noisy. To investigate this possibility, we asked whether the correlation structure differed between conditions where cycling started at the top of the cycle rather than at the bottom. The correlation structure was very similar in these two cases (compare *middle* and *left columns* of Figure 6E-F). This finding rules out the possibility that correlations are unstable simply because they are spurious, and demonstrates that not just any change in the task results in a change in the correlation structure. Changing the starting position had relatively little impact, while changing the performing arm had a dramatic impact.

Each matrix in Figure 6E-F corresponds to a given condition (a starting position and cycling direction). We wished to summarize, across all such conditions, the degree to which correlations are or aren’t preserved when the task is performed with one arm versus the other. To do so, for each condition and each pair of neurons we plotted their firing-rate correlation when the non-driven arm performed the task versus their correlation when the driven arm performed the task (Figure 6G). This is equivalent to plotting the values of the non-driven-arm correlation matrix (*right column* of Figure 6E,F) versus the corresponding values of the driven-arm correlation matrix (*left column*). Preserved correlations would yield diagonal structure. In fact, there was little tendency for correlated neurons to remain correlated, or for anti-correlated neurons to remain anti-correlated. The ‘meta-correlation’ was 0.1 and 0.05 (monkey E and F). Thus, if two neurons responded similarly when the driven arm performed the task, this said little regarding whether those neurons would respond similarly when the non-driven arm performed the task.

### Population activity is isomorphic across hemispheres

The above results demonstrate that both individual-neuron responses and their correlation structure are very different depending on which arm is employed to perform the task. One potential explanation is that very different signals are present: perhaps muscle-like signals when employing the driven arm versus more abstract signals when employing the non-driven arm. An alternative explanation is that many of the same signals are present, yet are reflected differently at the level of individual neurons. We have argued that motor cortex carries both muscle-like signals and non-muscle-like signals fundamental to the underlying computations (Churchland et al., 2012; Russo et al., 2018). However, those experiments examined only the driving cortex; it remains unknown which signals are shared with the non-driving motor cortex.

We first asked whether muscle-like signals are present in both hemispheres. We trained a regularized linear decoder to predict performing-arm muscle activity based on neural activity. We assessed generalization to a held-out condition, repeating this procedure for each condition. Both the driving and non-driving cortex accurately predicted muscle activity (Figure 7A). Across all conditions, generalization *R*^*2*^ was high for both the driving and non-cortex (Figure 7B, generalization performance computed across all muscles). Generalization performance was lower for the non-driving cortex, but this was a small effect and was significant for only one monkey (p= 0.15 and p = 0.016 for monkey E and F, two-sided Wilcoxon signed rank test across 8 conditions). We thus saw no evidence that signals related to muscle activity were absent in the non-driving cortex.

**Figure 7:**
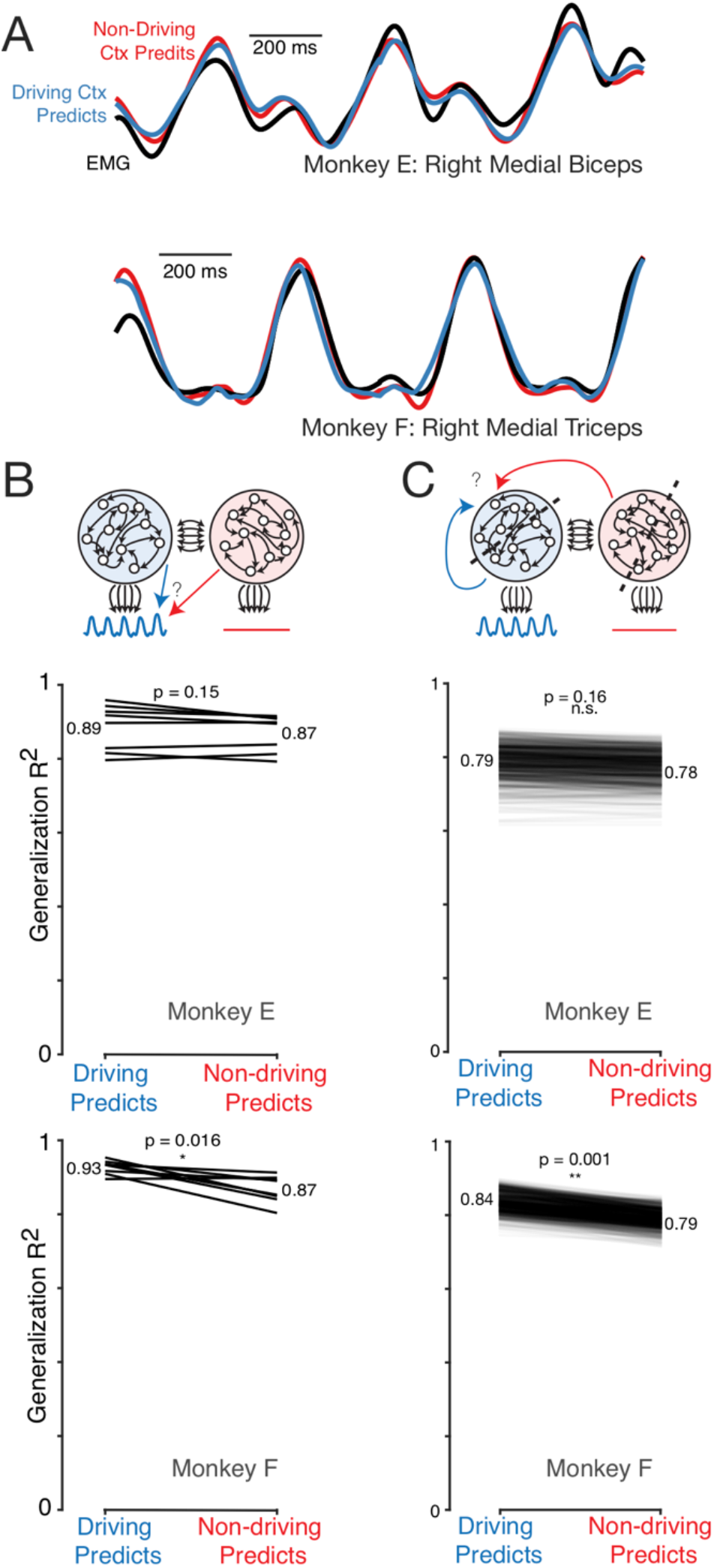
Population activity in the driving and non-driving hemispheres carries similar signals. (A) Muscle activity (*black*) and predictions of muscle activity based on a linear decode of neural activity in the driving (*blue*) and non-driving (*red*) cortex. Examples are shown for two muscles from the two monkeys. In both cases, data is from a test condition and illustrates generalization performance. (B) Quantitative comparison of performance when predicting muscle activity from neural activity in the driving versus non-driving hemisphere. *Top*: Cartoon illustration of the analysis approach. Performing-arm muscle activity (*blue trace*) is a product of descending connections (*black arrows*) from neurons within the driving cortex (*blue circle*). It should thus be possible to predict (*blue arrow*) that muscle activity from neural activity recorded from the driving cortex. The presence or absence of muscle-like signals within the non-driving cortex (red circle) was assessed by asking how well such activity predicted (*red arrow*) performing-arm muscle activity. *Bottom two panels*: prediction performance for the above comparisons, for both monkeys. Each line corresponds to one behavioral condition, and shows the percent variance (of muscle population activity) predicted by the driving and non-driving cortices. (C) Analysis asking whether the signals carried by the driving cortex are also present in the non-driving cortex. *Top*: Cartoon illustration of the analysis approach. When the task is performed with a given arm, neural activity within the corresponding driving cortex is predicted either from the activity of other neurons either within the driving cortex (blue arrow) or within the non-driving cortex (red arrow). *Bottom two panels*: Prediction performance for those two comparisons, for both monkeys. Each line shows the performance for one random split of the data. After each random split, the activity of half the neurons in the driving cortex was predicted either based on the other half, or based on a matched number of neurons from the non-driving cortex.

Might other signals be restricted to the driving cortex? Rather than attempting to infer specific candidate signals, we developed a method to address this question generically (Figure 7C, top). We randomly divided driving-cortex neurons into two groups. Because the division is random and the number of neurons large, each group should reflect approximately the same set of signals. Thus, one group should be able to accurately predict activity in the other group. This was indeed the case (Figure 7C, ‘Driving Predicts’). We next generated a size-matched group of neurons from the non-driving cortex, and asked whether their activity could be used to predict activity in the driving cortex. If all signals are present in both hemispheres, it should be possible to predict driving-cortex activity based on non-driving cortex activity. Conversely, if a major signal is missing in the non-driving hemisphere, prediction would be compromised.

Driving cortex activity was predicted almost as well, based on activity in the opposite hemisphere, as it had been based on activity within the same hemisphere (Figure 7C, thin lines show results for 1000 random divisions). For Monkey E, the difference was non-significant (p = 0.16, bootstrap test across 1000 resamples). For Monkey F, the difference was significant but small: a loss of 5% of the variance explained (p = 0.001, bootstrap across 1000 resamples). Thus, we saw no evidence for large signals that are present in the driving cortex but absent in the non-driving cortex.

### Neural trajectory tangling is low for both hemispheres

We recently described a major difference between M1 population activity and downstream muscle activity (Russo et al., 2018). Only M1 avoids ‘trajectory tangling,’ defined as the occurrence of similar population states with very different derivatives. Trajectory tangling becomes high if the population trajectory crosses itself, or if the trajectory for one condition traverses near that for another condition but travels in a different direction. Pattern-generating recurrent networks are noise-robust only if trajectory tangling is low, suggesting an explanation for why low trajectory tangling was observed in M1. It is unknown whether the non-driving cortex participates (via callosal connections) in pattern generation. We therefore wondered whether the non-driving cortex would show similarly low trajectory tangling. Notably, it is possible for a cortical area to be active during cycling yet have high trajectory tangling; this was true of proprioceptive primary somatosensory cortex (Russo et al., 2018).

We computed the tangling index (as used in Russo et al. 2018) for every time during the middle cycles, across all conditions. We did so for population activity in the driving and non-driving cortex, and also for the muscle populations, and compared the resulting distributions. The muscles often showed high trajectory tangling, revealed by a long right tail in the cumulative distributions (Figure 8A,B, *black lines*). The driving cortex displayed consistently low trajectory tangling: cumulative distributions (*blue*) plateaued early. This replicates prior results: trajectory tangling is much lower for the driving cortex than for the downstream muscle population. Notably, this is true even though single-neuron and single-muscle responses are superficially similar.

**Figure 8:**
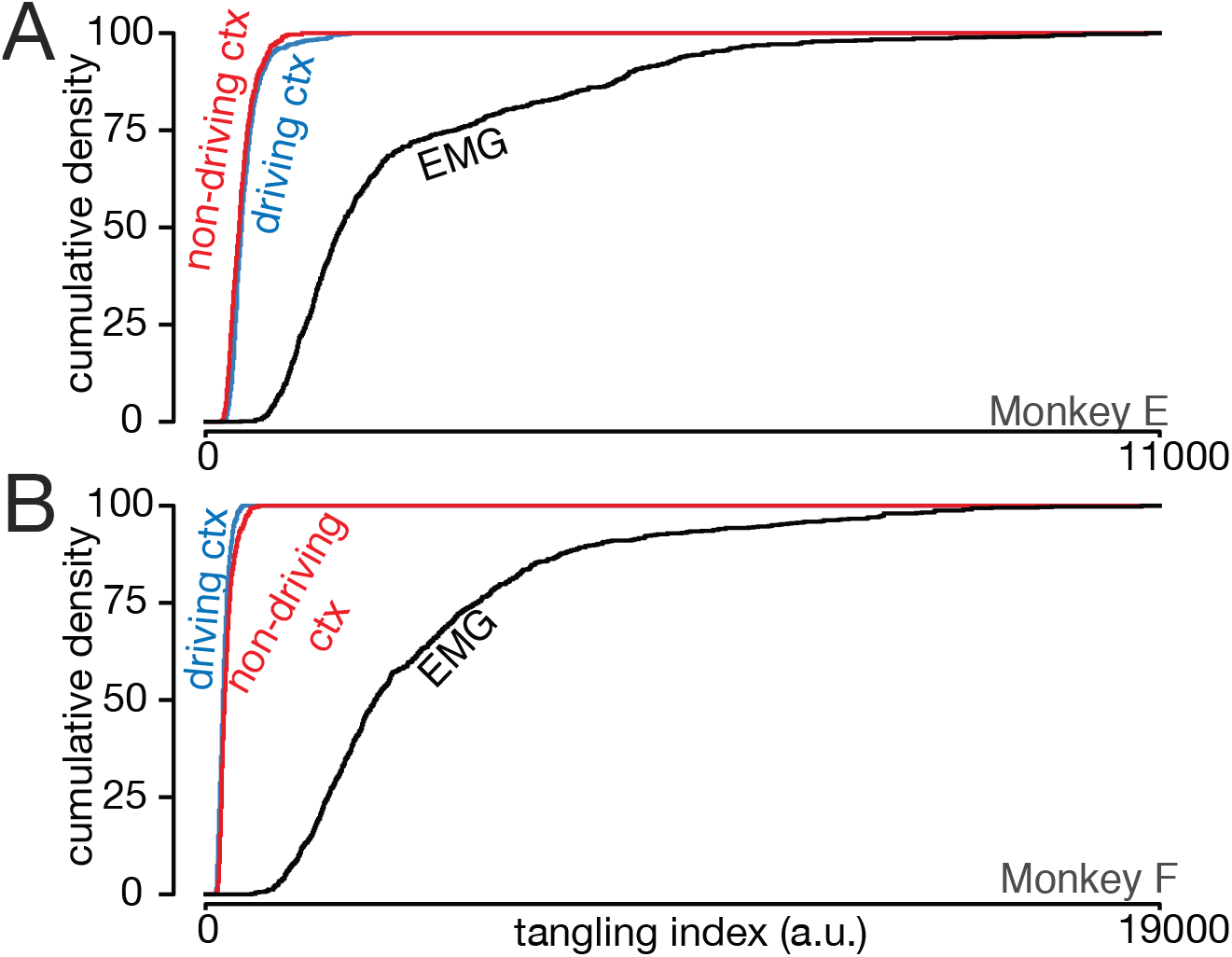
Trajectory tangling is similar for the driving and non-driving cortices. (A) Cumulative distribution of trajectory tangling for the driving cortex (*blue*), non-driving cortex (*red*), and muscle activity (*black*). Distributions were calculated across all time points and conditions. Data for monkey E. (B) As in A, for Monkey F.

Trajectory tangling was also low for the non-driving cortex (*red*). For monkey E, tangling was slightly higher in the non-driving versus driving cortex (468 ± 201 versus 420 ± 153; mean ± S.D.) while the opposite was true for monkey F (374 ± 105 versus 430 ± 146). Thus, trajectory tangling is similarly low for both cortices, with only small and inconsistent differences. Critically, for both the driving and non-driving cortex, neural trajectory tangling was much lower than muscle trajectory tangling. The latter averaged 2296 ± 1766 for Monkey E and 4392 ± 2950 for Monkey F. Taken together with the results above, we found little hemispheric difference regarding either the major signals or the organization of population trajectories.

### Neural activity occupies different dimensions for movements of different arms

If similar signals are present regardless of which arm moves, how does the brain avoid moving the wrong arm? In confronting this question, we took inspiration from recent work suggesting that only some neural dimensions in motor cortex are ‘muscle potent’; activity in those dimensions produces output that will influence the muscles. Other dimensions are ‘muscle null’; activity in those dimensions has no direct outgoing impact on muscle activity (Druckmann and Chklovskii, 2012; Kaufman et al., 2014). The presence of output-null dimensions is natural (and typically inevitable) when patterns are generated by a recurrent network with more internally-connected neurons than output neurons. We wondered whether this principle might apply to the present case. We considered all recorded neurons, across both hemispheres, as a unified population. We asked whether signals related to the movement of each arm are partitioned in a manner that could allow signals related to one arm to naturally avoid impacting the other arm.

We used principal component analysis (PCA) to find neural dimensions that best explain activity. We applied PCA once for conditions where the right arm performed the task (‘right-performing’ conditions) and again for conditions where the left arm performed the task (‘left-performing’ conditions). A ‘right-arm’ space was defined by the PCs found for the right-performing conditions. A ‘left-arm’ space was defined analogously. We were interested in what occurred in the right-arm space when the left arm performed the task, and vice versa. Importantly, in both cases PCA considered the responses of the same unified population of neurons (all recorded neurons across both hemispheres).

The right-arm space captured (by construction) population activity when the right arm performed the task. This can be appreciated in Figure 9A,C by the large near-circular trajectories for the right-performing conditions (*red*). The rapid rise in cumulative variance accounted for (Figure 9B,D, *red*) reveals that a small number of right-arm PCs successfully captured most of the variance for the right-performing conditions. In contrast, the right-arm space did not effectively capture variance for the left-performing conditions. Left-performing neural trajectories are small when projected onto the right-arm PCs, with little clear structure (Figure 9A,C, *blue*). The cumulative percent variance accounted for (left-performing conditions projected onto the right-arm PCs) rose slowly (Figure 9B,D, *blue*). Analogous results were found when analyzing the left-arm space (Figure 9E-H).

**Figure 9:**
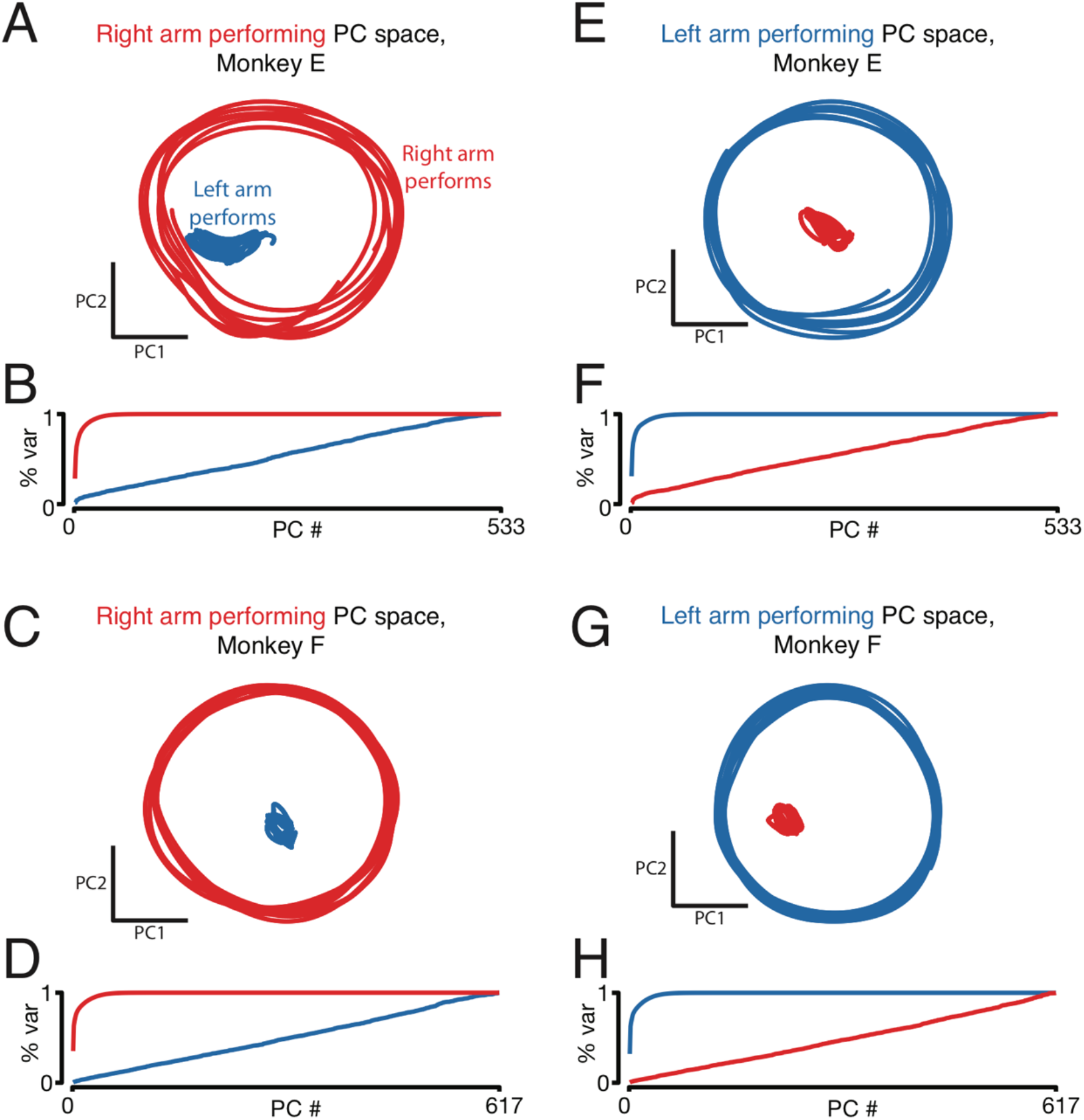
Right-arm-related and left-arm-related population activity lie in orthogonal subspaces. (A) Projection of population activity for right-arm-performing (*red*) and left-arm-performing (*blue*) conditions, with PCs found using only right-arm-performing data. The neural population includes all recorded neurons across both hemispheres. Analysis considers data from the middle cycles when cycling forward, for both top-start and bottom-start conditions. Data are for monkey E. (B) Cumulative variance explained for right-arm-performing (*red*) and left-arm-performing (*blue*) conditions, using PCs found from right-arm-performing data only. (C,D) As above but for Monkey F. (E-H) As in A-D, but with PCs found using only left-arm-performing data. All data shown are for analyses performed on forward cycling conditions.

Thus, relatively little variance was captured when activity for left-performing conditions was projected onto the right-arm space, and vice versa. Averaged across all such conditions, the top 5 PCs explained 7 ± 1% (monkey E) and 2 ± 0.5% (monkey F) of the variance (mean ± std. computed across conditions). This was in contrast to the large amount of variance captured when conditions were projected onto the top 5 PCs of their ‘own’ space: 80 ± 2% (monkey E) and 80 ± 2% (monkey F). This asymmetry was not simply due to projecting onto a space built from the same data versus other data. For example, the top 5 PCs based on top-start conditions captured almost as much variance during bottom-start conditions (63 ± 4% for monkey E and 66 ± 2% for monkey F) as during top-start conditions (79 ± 2% and 79 +/− 2%).

Thus, neural responses related to the two arms occupy nearly orthogonal subspaces. Dimensions that robustly capture activity when one arm performs the task do not continue to do so when the other arm performs the task. This could occur trivially if neurons fall into two groups: neurons that respond only when the right arm performs the task, and neurons that perform only when the left arm performs the task. However, as documented above, no such separation was present. The distribution of arm preference indices was unimodal with a median near zero (Figure 4B) indicating that most neurons responded regardless of which arm performed the task. The orthogonality of dimensions is instead related to the finding that the correlation structure depends strongly on which arm performs the task (Figure 6). As such, this is intrinsically a population-level finding that could not have been inferred from analyses focused on individual neurons.

### Linear decoders naturally separate signals related to the two arms

The separation of activity into orthogonal subspaces may allow descending control of one arm to naturally ignore signals related to the other arm. To test the plausibility of this hypothesis, we trained linear decoders to predict, based on the activity of the entire neural population, muscle activity for a given arm. The decoder was trained only using conditions where that arm performed the task. For example, the decoder was trained to predict muscle activity in the right arm while the right arm performed the task. Restricted to this situation, decoders performed well, predicting a median of 91% (Monkey E) and 93% (Monkey F) of the variance on held-out conditions. Examples of predicted muscle activity (Figure 10A,D, *orange traces at top*) are shown for one muscle for each monkey. We then assessed generalization to conditions where the other arm performed the task. For example, a decoder that fit right-arm muscle activity (trained only on right-performing conditions) was asked to generalize and predict right-arm muscle activity during left-performing conditions. Does the prediction stay relatively flat, accurately capturing the absence of muscle activity? Or does the decoder become contaminated by signals related to the performing arm?

**Figure 10:**
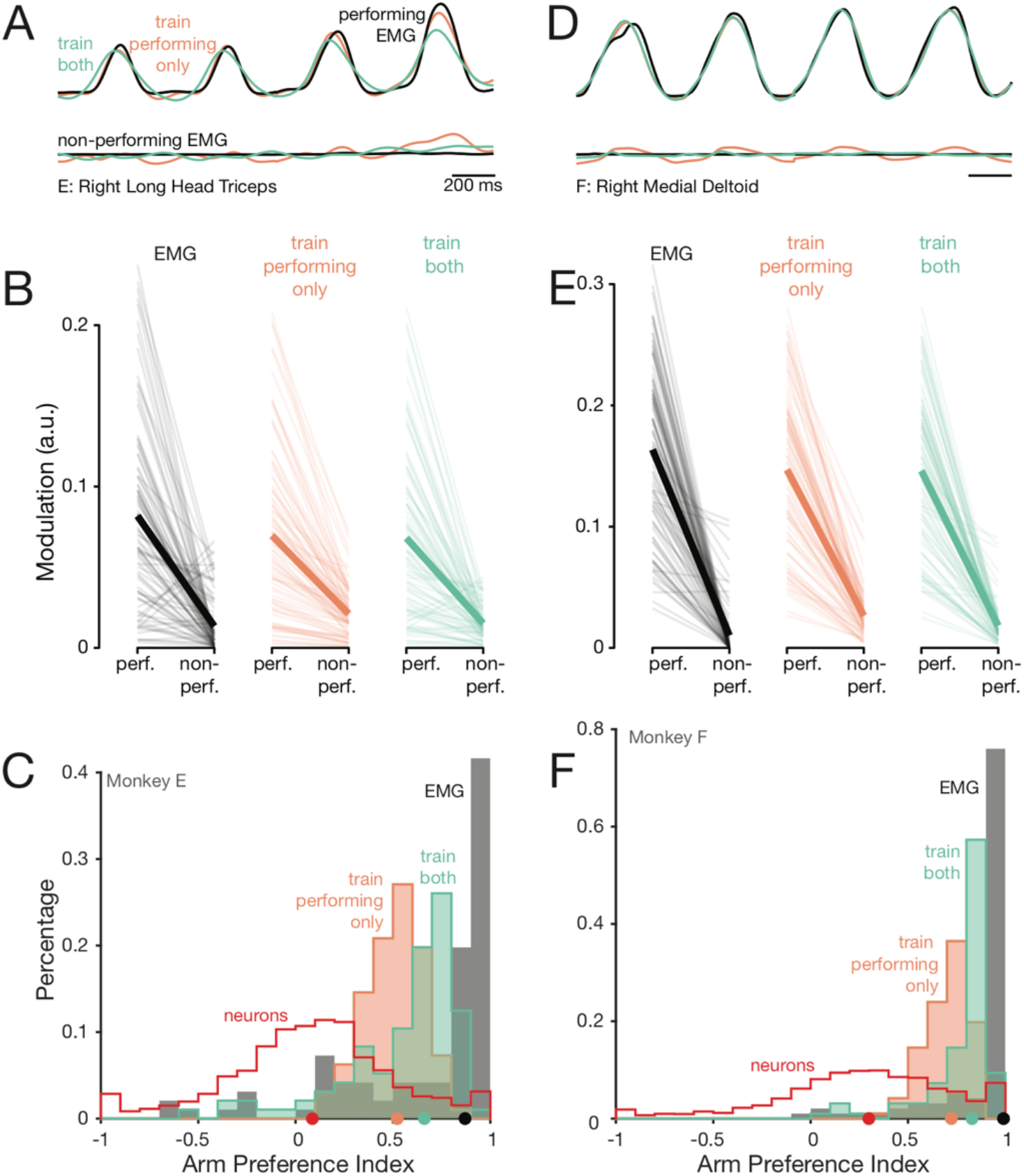
When decoding muscle activity in a given arm, decoders naturally ignore activity related to the other arm. (A) Activity of the long head of the *triceps*, recorded from the right arm of monkey E. *Black trace:* Recorded EMG activity. *Orange trace*: Prediction of a decoder trained only on right-arm-performing conditions. *Green trace*: Prediction of a decoder trained on a subset of both left- and right-arm-performing conditions. Top traces: while the right arm performed the task. Bottom traces: while the other arm performed the task. Data are for a left-out condition, to which decoders had to generalize. (B) For Monkey E, EMG modulation while performing the task versus while the other arm performs the task. *Thin lines*: individual muscles and conditions. *Thick lines*: median modulation. *Black*: Recorded EMG activity. *Orange:* Predictions of a decoder trained only on performing conditions. *Green*: Predictions of a decoder trained on both performing and non-performing conditions. (C) For Monkey E, distribution of arm preference indices for neurons (*red)*, muscles (*gray*), predictions of a decoder trained only on performing-arm conditions (*orange*), predictions of a decoder trained on both performing and non-performing arm conditions (*green*). *Red histograms* and *gray histograms* differ slightly from those in Fig. 4A-B as they are computed per condition, given that the present analysis focuses on generalization performance for left-out conditions. (D-E) As in A-C, for Monkey F. All data shown is from generalization to held-out conditions.

Decoders accurately generalized, and predicted little modulation of muscle activity in the non-performing arm. For the two example muscles shown, the predicted muscle activity was relatively flat, in agreement with the lack of modulation of the empirical muscle activity (Figure 10A,D, *bottom*, compare *orange* and *black traces*). This was true across muscles and conditions: decoded muscle activity was only weakly modulated for conditions when the task was performed with the other arm (Figure 10 B,E, *orange*), even though the decoder was not trained on such conditions and even though the neural activity upon which the decoder was based was similarly modulated regardless of the performing arm. The ability of the decoder to ignore such activity was inherited from the orthogonality of subspaces described above. When trained using right-arm conditions, decoders naturally employ right-arm dimensions. Because those dimensions are largely unoccupied when the left arm performs the task, the decode shows minimal modulation.

Due to these properties, decoders naturally produce predicted muscle activity with positive arm-preference indices (Figure 10C, *orange histograms*). These distributions are right-shifted relative to those for the neural activity upon which decoding was based (*red histogram)*. Thus, the structure of population activity ensures that a decoder, trained to extract activity related to one arm, will naturally tend to ignore activity related to the other arm.

Yet while decoders tended to naturally ignore activity related to the ‘wrong’ arm, this feature was imperfect: small amounts of residual modulation were still present (Figure 10A,D, orange trace at bottom) leading to arm-preference indices smaller than those of the muscles. Of course, one would expect improved ability to segregate activity if a decoder is trained to do so: i.e., to predict muscle activity both when the muscle is strongly modulated (when the relevant arm performs the task) and also when it is not (when the other arm performs the task). This was indeed the case (Figure 10, *green*). Note that these decoders still had to generalize to left-out conditions; they simply had the benefit of training data that included both left- and right-performing conditions.

There is thus no paradox in the absence of muscle activity in the non-performing arm, despite robust neural activity across both hemispheres. Signals related to the two arms are separated into different neural subspaces. As a result, even simple linear decoders naturally separate signals related to one arm versus the other.

## Discussion

We found that neural signals related to movements of the right and left arms are anatomically intertwined. Signals were mixed across hemispheres; when one arm moved, neurons in both hemispheres were modulated. Signals were also mixed across neurons; most neurons responded when both their driven and their non-driven arm performed the task. Individual neurons responded very differently depending on which arm was moving. Yet at the level of the population, both hemispheres contained similar information. Surprisingly, we did not find signals that were strongly present in the driving cortex but absent in the non-driving cortex. This was true even for muscle-like signals, which could be decoded similarly well from either hemisphere.

Despite this intermixing, signals corresponding to the two arms were highly separable at the level of neural dimensions. Activity related to the left arm occupied a set of dimensions nearly orthogonal to the dimensions occupied by activity related to the right arm. As a result, even simple linear decoders could read out commands for one arm while ignoring commands for the other arm.

### Separation of information across dimensions is a common feature of cortical activity

Our results contribute to an increasingly broad set of studies reporting that neural activity related to different computations or task parameters is often separated across neural dimensions, instead of at the level of brain areas or individual neurons. During reaching, dimensions carrying preparatory activity are orthogonal to dimensions carrying muscle-related signals (Kaufman et al., 2014), and more generally to all dimensions occupied by movement-related activity (Elsayed et al., 2016). Activity related to the timing of movement initiation and activity related to which movement is being generated also occupy orthogonal dimensions (Kaufman et al., 2016). In sensory decision making, different aspects of the task (e.g., stimulus versus task time, color versus direction of the stimulus, or auditory versus visual cues) are integrated in different neural dimensions in prefrontal cortex (Machens et al., 2010; Mante et al., 2013) and parietal cortex (Raposo et al., 2014). Separating neural activity into separate dimensions for separate computations may thus be a general strategy used by many brain regions. A potential advantage of this strategy is that signals are able to interact (e.g., in the case where movements of two limbs are coordinated) yet can still be read out separately (Druckmann and Chklovskii, 2012; Kaufman et al., 2014)

### Comparison with prior studies of lateralization in M1

Our finding that individual neurons often respond during movements of either arm is in broad agreement with prior primate recording studies. A majority of these studies describe intermixing of right- and left-arm responses in the activity of individual M1 neurons (Cisek et al., 2003; Donchin et al., 2002, 1998; Kermadi et al., 1998; Steinberg et al., 2002). However, a handful of these studies report a much smaller percentage of ipsilateral (non-driving) neural responses (Aizawa et al., 1990; Tanji et al., 1988). These studies examined neural responses in the hand area of M1 during small finger movements. The hand area of M1 has fewer callosal connections and fewer ipsilateral projections than the arm area of M1 has (Jenny, 1979; Jones and Wise, 1977; Rouiller et al., 1994). Thus, a likely explanation for varied prior results is that hand-related neural computations are more divided by hemisphere, while arm-related computations are largely intermixed. An alternative explanation is that it is simply easier to move one hand while keeping the other still, resulting in greater neural segregation due to better experimental control over lateralization of muscle activity. Our results support the first explanation; muscle activity was weak in the non-driven arm, and the very small movements of that arm had no statistical impact on neural activity.

Of prior studies, two explicitly compared neural response properties (e.g., preferred directions) during unimanual movements of each arm. Cisek et al. (2003) found that neurons in M1 had limb-dependent preferred directions, yet Steinberg et al. (2002) reported preserved preferred directions. At the same time, Steinberg et al. found that left-arm and right-arm reach directions could be independently decoded by separate pools of neurons, suggesting some degree of limb-dependence. The present findings indicate that neural responses are strongly limb-dependent. Responses during performance with one versus the other arm were weakly correlated at the single-neuron level, and occupied nearly orthogonal subspaces at the population level.

### Possible reasons for ipsilateral motor cortical activity

There exist multiple reasons why motor cortex might be active when the non-driving arm performs the task. A straightforward possibility is that motor cortex employs an abstract limb-independent representation of movement. However, this hypothesis is unlikely given the strongly limb-dependent nature of responses. Alternatively, the two cortices may process different but complementary information. This hypothesis is also unlikely; we found no large signals that were present in the driving cortex but absent in the non-driving cortex.

It is also possible that activity ipsilateral to the moving arm may relate to uncrossed descending connections (Kuypers, 1981; Rosenzweig et al., 2009). For example, activity in the right motor cortex could exist to drive, via uncrossed connections, muscle activity in the right arm when that arm performs the task. Our results are in principle consistent with this hypothesis. However, prior studies have found little evidence for a robust relationship between M1 and ipsilateral muscle activations. Intracortical microstimulation readily produces contralateral muscle responses (Kwan et al., 1978; Sessle and Wiesendanger, 1982), yet very rarely generates ipsilateral muscle responses (Aizawa et al., 1990). Intracellular recordings of motoneurons reveal no monosynaptic evoked potentials from ipsilateral corticospinal tract stimulation and spike-triggered EMG effects are present only for contralateral muscles (Soteropoulos et al., 2011). For these reasons, we suspect that uncrossed projections are unlikely to be the primary reason that the non-driving motor cortex is active.

Another possibility is that activity in the non-driving cortex is produced by an efference copy of signals generated and employed by the driving cortex, which are conveyed to allow coordination between the limbs. Many – perhaps most – movements require coordination across the midline. Given the near-ubiquitous need for coordination, it may be that efference copy signals are simply conveyed by default, and ignored if they are not needed. Our results are consistent with this possibility, and argue that if it is correct, then the relevant efference copy must be quite complete. That is, the driving cortex must convey the majority of the signals it generates, rather than (for example) just the output signals.

A final, and intriguing possibility, is that motor cortical computations are largely distributed across both hemispheres (Li et al., 2016). In the extreme, neurons in the non-driving cortex might simply be viewed the way we view most neurons in the driving cortex; they can contribute to the computation even if they are one or more synapses from the cortico-spinal neurons that will convey the output. This hypothesis is appealing because it could explain the finding that all major signals appear to be shared between hemispheres. More generally, if a randomly chosen neuron from the non-driving cortex has responses that are nearly indistinguishable from a neuron chosen from the driving cortex, perhaps our default assumption should be that they are participating in the same computation. While appealing, it is unclear if this hypothesis can be reconciled with the finding that motor cortex inactivation principally affects the contralateral limbs (Glees and Cole, 1950; Liu and Rouiller, 1999; Passingham et al., 1983). If both hemispheres participate in controlling both arms, one would expect a more bilateral deficit. A possible, but highly speculative, resolution is that the network is sufficiently robust that it can still function when many neurons are inactivated, so long as the output neurons can still convey the necessary commands.

### Ipsilateral arm signals in other brain regions

Cortical areas, subcortical areas, and the spinal cord all contribute to the control of dexterous movements. Indeed, other studies comparing contralateral and ipsilateral movements have found that not only M1, but also the dorsal premotor cortex (Cisek et al., 2003; Kermadi et al., 2000; Tanji et al., 1988), ventral premotor cortex (Michaels and Scherberger, 2018), Supplemental Motor Area (Donchin et al., 2002; Gribova et al., 2002; Kazennikov et al., 1999; Kermadi et al., 2000, 1998; Tanji et al., 1988), Cingulate Motor Area (Kermadi et al., 2000), and the Posterior Parietal Cortex (Kermadi et al., 2000) all contain neurons which respond to movements of the ipsilateral arm. Furthermore, there are circuits in the brainstem and spinal cord which specifically support the generation of coordinated, rhythmic movements like locomotion (Duysens and de Crommert, 1998). Other brain regions, such as the Anterior Intraparietal Area, encode movement parameters in a largely effector-independent manner (Michaels and Scherberger, 2018), suggesting that the relationship of motor and visuo-motor areas to ipsilateral movements may vary depending on their role in motor computation. In general, movement generation is the result of the action of a broad, interconnected network of brain and spinal regions. We focused on M1 because it is the cortical region that, based on anatomy and microstimulation results, seemed most likely to have a lateralized representation of movement. Yet even in M1 we failed to find evidence of strongly lateralized activity.

### Summary

Neural signals related to movements of either arm were mixed, both within hemispheres and within single neurons. However, signals related to the two arms were naturally partitioned into different neural dimensions. This underscores the computational usefulness of leveraging different dimensions for different computations; signals can be shared across a wide population yet still be readily separated by downstream regions. Our results argue that motor cortex shares highly detailed information cross-cortically, suggesting that control may span both hemispheres even if output commands originate primarily from the contralateral hemisphere.

## Acknowledgements

We thank Y. Pavlova for expert animal care. A. Russo provided code to calculate trajectory tangling. This work was supported by NINDS 1DP2NS083037, NIH CRCNS R01NS100066, NINDS 1U19NS104649, the Simons Foundation (SCGB#325233 and SCGB#542957), the Grossman Center for the Statistics of Mind, the McKnight Foundation, P30 EY019007, a Klingenstein-Simons Fellowship, the Searle Scholars Program. KCA was funded by a Simon’s Foundation Collaboration on the Global Brain Post-doctoral Fellowship.

